# Pyphe: A python toolbox for assessing microbial growth and cell viability in high-throughput colony screens

**DOI:** 10.1101/2020.01.22.915363

**Authors:** Stephan Kamrad, Maria Rodríguez-López, Cristina Cotobal, Clara Correia-Melo, Markus Ralser, Jürg Bähler

## Abstract

Microbial fitness screens are a key technique in functional genomics. We present an all-in-one solution, *pyphe*, for automating and improving data analysis pipelines associated with large-scale fitness screens, including image acquisition and quantification, data normalisation, and statistical analysis. *Pyphe* is versatile and processes fitness data from colony sizes, viability scores from phloxine B staining or colony growth curves, all obtained with inexpensive transilluminating flatbed scanners. We apply *pyphe* to show that the fitness information contained in late endpoint measurements of colony sizes is similar to maximum growth slopes from time series. We phenotype gene-deletion strains of fission yeast in 59,350 individual fitness assays in 70 conditions, revealing that colony size and viability provide complementary, independent information. Viability scores obtained from quantifying the redness of phloxine-stained colonies accurately reflect the fraction of live cells within colonies. *Pyphe* is user-friendly, open-source and fully-documented, illustrated by applications to diverse fitness analysis scenarios.

## Introduction

Colony fitness screens are a key technique in microbial genetics. The availability of knock-out libraries has revolutionised forward genetics (Giaever and Nislow, 2014), and large collections of wild isolates (Jeffares et al., 2015; Peter et al., 2018) as well as synthetic populations (Bloom et al., 2013; Cubillos et al., 2013) have proven a powerful tool to study complex traits. More recently, high-throughput phenomics, the systematic measurement of fitness for hundreds of conditions and/or hundreds/thousands of strains in parallel, is driving our systems-level understanding of gene function (Brochado et al., 2018; Costanzo et al., 2016; Kuzmin et al., 2018; Nichols et al., 2011).

Screens generally follow a workflow where strains are arranged in high-density arrays (e.g. 384 or 1536 colonies per plate) and transferred using a colony-pinning robot or manual replicator. Image analysis software enables fast and precise quantification of colony sizes and other phenotypes (Bischof et al., 2016; Kritikos et al., 2017; Lawless et al., 2010; Memarian et al., 2007; Wagih and Parts, 2014). Colony-size data is prone to noise and technical variation between areas on the same plate and across plates and batches, some of which can be corrected by normalisation procedures (Baryshnikova et al., 2010; Blomberg, 2011; Zackrisson et al., 2016). Finally, differential fitness is assessed statistically, for which specialised approaches are available (Collins et al., 2010, 2006; Wagih and Parts, 2015).

Most screens use a single image or time point per plate (an endpoint measurement). Potentially more information is contained in the growth of colony sizes over time, and a low-resolution time course of colony sizes can be used to fit growth models to population size data (Addinall et al., 2011; Banks et al., 2012; Shah et al., 2007). High-resolution image time series contain potentially even more information and have been used to determine lag phases (Levin-Reisman et al., 2014). Recently, very precise fitness determination has been achieved by high-resolution, transilluminating time course imaging and growth curve analysis, using the maximum slope of the obtained growth curve and reference grid normalisation (Zackrisson et al., 2016). The parallel use of commercially available scanners, combined with high-density arrays of colonies can enable phenotyping at very large scales, but poses challenges in terms of data storage, processing, equipment and the need for temperature-controlled space.

The dead-cell stain phloxine B can provide an additional phenotypic readout related to the proportion of dead cells in a colony. Phloxine B has previously been used to assess the viability of cells in budding yeast by microscopy (Tsukada and Ohsumi, 1993) and in fission yeast colonies (Matynia et al., 1998). When applied in a screening context, colonies are assigned a score which reflects the ‘redness’ of the colony to serve as an additional quantitative phenotype that can be used for downstream analysis (Lie et al., 2018).

Despite the popularity and importance of microbial colony screens, a consensus data framework has so far not emerged. In our laboratories, fitness screens are an essential technique used on a variety of scales, from a handful of plates to several thousands, and by researchers with varying bioinformatics skills. To enable and standardise data analysis workflows, we have developed a bioinformatics toolbox with a focus on being versatile, modular and user friendly. *Pyphe* (*py*thon package for *phe*notype analysis) consists of 5 command line tools each performing a different workflow step as well as the underlying functions provided as a python package to expert users.

We illustrate the use of *pyphe* by investigating the growth dynamics of 57 natural *S. pombe* isolates. We show that the spatial correction implemented in *pyphe*, based on that proposed by (Zackrisson et al., 2016), is effective in reducing measurement noise without overcorrection. Late endpoint measurements are shown to provide similar readouts to maximum slopes, but with lower precision. We then investigate the relationship between colony sizes and viability scores in a broad panel of *S. pombe* knock-out strains in over 40 conditions and find that the two approaches provide orthogonal and independent information. Using imaging flow cytometry, we link colony redness scores to the percentage of dead cells in a colony and show that phloxine B staining provides similar results as a different live/dead stain.

## Results

### *Pyphe* enables analysis pipelines for fitness-screen data

The *pyphe* pipeline (Fig. 1) is designed to take different fitness proxies as input: endpoint colony sizes, colony growth curves, or endpoint colony viability estimates from phloxine B staining. Image acquisition, image analysis, growth-curve analysis, data normalisation and statistical analysis are split into separate tools which can be assembled into a pipeline as required for each experiment and combined with other published tools, e.g. *gitter (Wagih and Parts, 2014)* for image analysis. Each tool takes and produces human-readable data in text/table format.

**Figure 1:**
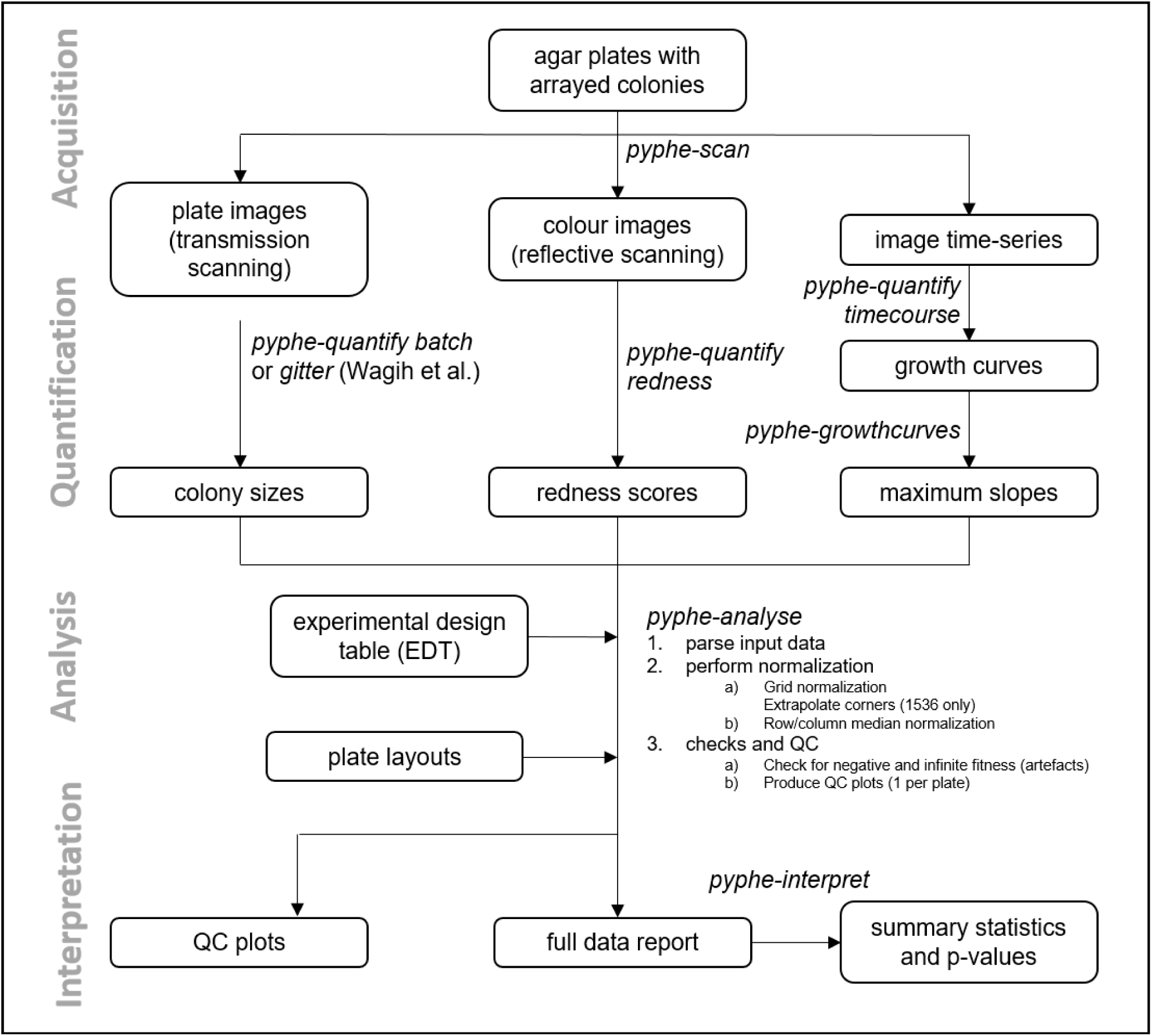
Data processing workflows using *Pyphe*. *Pyphe* is flexible and can use several fitness proxies as input. In a typical endpoint experiment, plate images are acquired using transmission scanning and colony sizes are extracted using the R package *gitter (Wagih and Parts, 2014)*. Alternatively or additionally, plates containing phloxine B are scanned using reflective scanning and analysed with *pyphe-quantify* in redness mode to obtain redness scores reflecting colony viability. Alternatively, image time series can be analysed with *pyphe-quantify* in *timecourse* mode and growth curve characteristics extracted with *pyphe-growthcurves. Pyphe-analyse* analyses and organises data for collections of plates. It requires an Experimental Design Table (EDT) containing a single line per plate and the path to the data file, optionally the path to the layout file, and any additional metadata the user wishes to include. Data is then loaded and the chosen normalisation procedures are performed. QC plots are produced and the entire experiment data is summarised in a single long table. This table is used by *pyphe-interpret* which produces a table of summary statistics and p-values for differential fitness analysis.

In a typical workflow, images are acquired using *pyphe-scan* which provides an interface for image acquisition using SANE (Scanner Access Now Easy) on a unix-type operating system. It handles plate numbering, cropping and flopping, and format conversion functionality for large image stacks. Optionally, image time-series can be recorded. *Pyphe-scan* is written to work with EPSON V800 scanners as established by (Zackrisson et al., 2016).

Colony properties are then quantified from images using *pyphe-quantify* which can operate in three different modes. In *batch* mode (for colony-size quantification using grayscale transmission scanning) or *redness* mode (colony-viability estimation using phloxine B and reflective colour scanning) it separately analyses all images that match the input pattern (by default all jpg images in working directory), producing a csv table and qc image for each. In *timecourse* mode, colony positions are determined in the last image and the mask applied to all previous images, extracting background-subtracted mean intensities for each colony/spot and producing a single table with the growth curves (one per column). It reports a wide range of colony properties: colony area, overall intensity (an estimator that reflects thickness as well as area), circularity, perimeter and centroid coordinates, making this tool useful in cases where colonies are not arrayed. The algorithm is described in detail in Supplementary Text 1 and Supplementary Fig. 1.

Spatial normalisation is performed for each plate, and data across all plates is aggregated using *pyphe-analyse* to produce a single table for downstream hit calling and further analysis. *Pyphe* implements a grid normalisation procedure based on the one previously described (Zackrisson et al., 2016) as well as row/column median normalisation. We have modified the placement of the grids in 1536 format (Supplementary Fig. 2) and implemented checks for missing colonies and normalisation artefacts. The main output is a single long table, containing one row per colony, with all position-, strain-, meta- and fitness-data as well as details about the normalisation. Algorithms are further described in Supplementary Text 2 and Supplementary Fig. 2.

Finally, differential fitness is assessed using *pyphe-interpret* which produces summary statistics and p-values based on the complete data report from *pyphe-analyse*. The user specifies the columns to use for grouping replicates, for performing the test against, and for fitness (the dependent variable of t-test). Usually, we group by the condition column, specify the control condition, and use corrected and checked colony sizes to test for condition-specific fitness differences for each condition and strain using the final relative fitness estimate post normalisation.

### Effective normalisation reduces noise and bias in data

*Pyphe* is designed to use different fitness proxies as input. In particular, it can use either maximum growth rates extracted from growth curves or endpoint colony size measurements. Previous studies have reported that information from growth curves are more precise (Zackrisson et al., 2016), but their acquisition requires substantially higher investment and produces large amounts of image data. While lower precision could be easily compensated by a higher number of replicates, growth curves provide the additional advantage that they capture the entire growth phase instead of a static snapshot. The results obtained in endpoint measurements might therefore depend on the time point used for the measurement. For example, the fitness of a strain with a long lag phase but high maximum growth rate may be underestimated if an early time point is chosen.

To assess the extent to which the choice of time point matters, we recorded image time series for 57 *S. pombe* wild strains growing in 1536 (spots per plate) format on rich YES medium in approximately 20 replicates. These strains are genotypically and phenotypically diverse and should display a broad range of growth characteristics (Jeffares et al., 2015). Colonies were imaged every 20 minutes for a period of 48 hrs (145 measurement points for each of the 1536 spots). Then, colony areas were extracted with *gitter (Wagih and Parts, 2014)*, and corrected colony sizes were computed for each image using the grid normalisation with subsequent row/column median normalisation implemented in *pyphe* and averaged for each strain (Fig. 2A). A correlation matrix of all timepoints showed near perfect correlation of timepoints with the 48 hour end point from 16 hours when the rapid growth phase for most strains came towards its end (Fig. 2Biii). Notably, all timepoints were correlated with the initial timepoint, albeit much lower, suggesting a significant bias introduced by the amount of initially deposited biomass. In our hands, this problem is more pronounced with wild strains than with knock-out collections as the former exhibit a variable degree of stickiness. However, we overcome this issue by reporting strain fitness as a ratio of growth in an assay condition relative to a control condition, in which case this bias is neutralised. Later timepoints generally showed a much better correlation with maximum growth rate compared to early ones or those taken when growth is most rapid (Fig. 2Bi+ii). We conclude that late timepoints should be chosen for endpoint measurements, when the readout is stable but nonetheless correlates well with the maximum growth rate.

**Figure 2:**
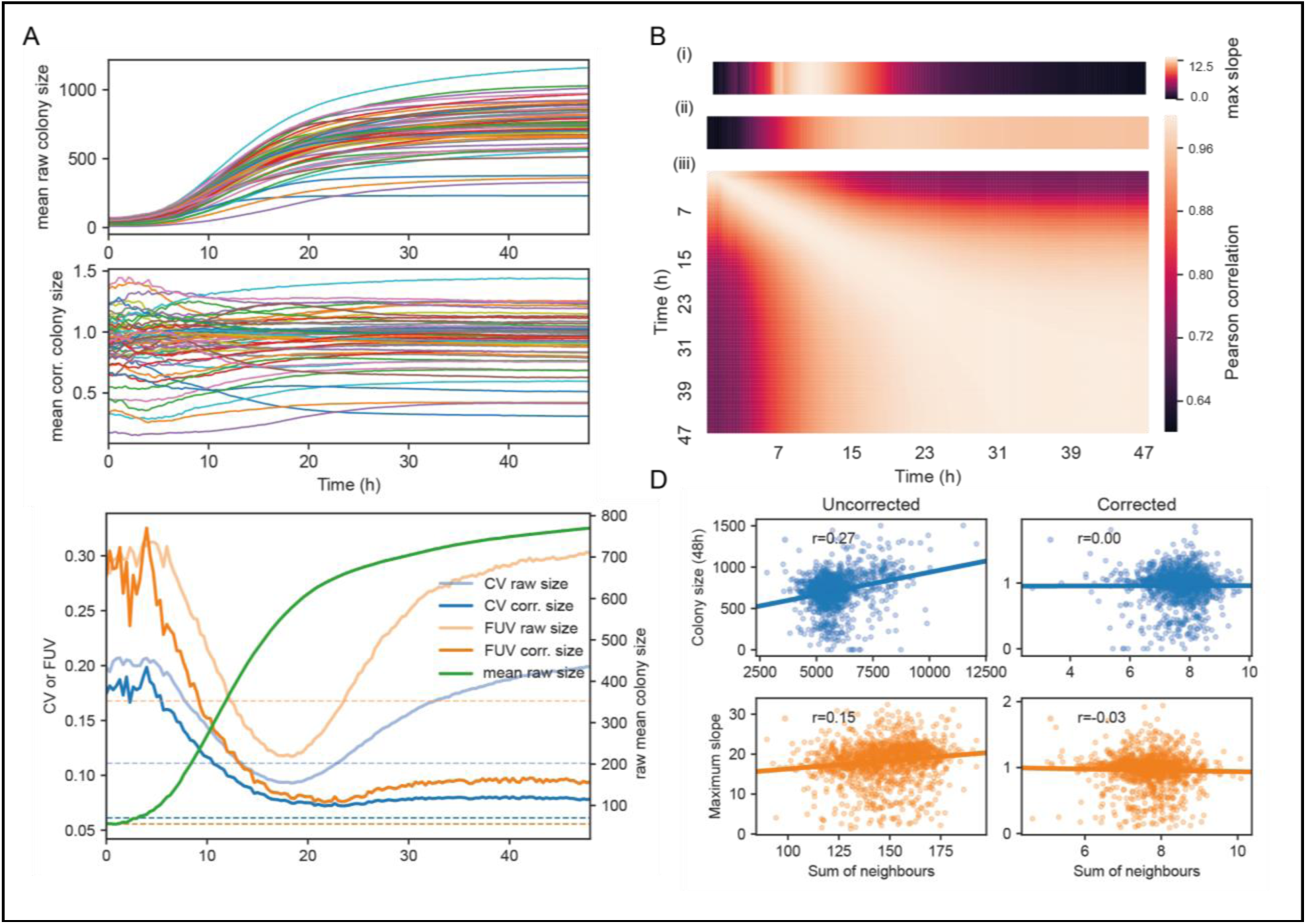
Normalisation strategies for growth curves and endpoints. **(A)** Growth curves of 57 wild *S. pombe* strains (average of 20 replicates each) before (top) and after (bottom) correction. Corrected colony sizes describe the fitness relative to the standard laboratory strain *972)*. **(B)** Late endpoint measurements are tightly correlated with maximum slopes. (i) Average growth rates (differences of uncorrected colony areas between consecutive timepoints, centered moving average with window size 3) across all strains. (ii) Pearson correlation of each individually corrected time point with corrected maximum slope of growth curves. The correlation increases throughout the rapid growth curve and then maintains high levels as the phase of fast growth comes to an end. (iii) Pearson correlation matrix of all corrected timepoints (averaged by strain prior to correlation analysis). **(C)** Coefficient of variation (CV, blue) and fraction of unexplained variance (FUV, orange) for corrected and uncorrected colony sizes throughout the growth curve. Dashed lines are the same values computed based on maximum slopes. The average growth curve of the control strain is shown in green. The normalisation procedure maintains noise at low levels even in later growth. Endpoint measurements contain slightly more noise than slope measurements. **(D)** Scatter plots of colony fitness estimates dependent on the sum of colony fitness of its 8 neighbours. A positive correlation, such as seen for the uncorrected readouts, points to spatial biases within plates (specific regions of a plate growing slower/faster, e.g. due to temperature, moisture or nutrient gradients). A negative correlation would be expected for competition effects. Without correction, regional plate effects dominate over competition effects and these are efficiently removed during grid correction. Importantly, the correction does not result in a negative correlation, a potential side-effect of correcting colony sizes by comparing it to the size of neighbouring controls, which would lead to phenotypes becoming artificially more extreme.

The choice of time point also affects the level of noise. The coefficient of variation (CV, the ratio of the standard deviation to the mean) of 112 replicates of the control strain dropped steadily during the rapid growth phase, reaching a minimum around 20 hours when it started to rise again (Fig. 2C). This is likely due to edge and other spatial effects which affect later growth as nutrients deplete and plates start to dry unevenly. After normalisation, the CV was generally lower, and this later rise in noise could be compensated so that the CV remained near its minimum. The CV of the maximum slopes was lower than obtained with endpoints. However, CV values alone are insufficient to judge the effectiveness of a normalisation strategy, as it reflects precision of the reported values but not their accuracy. As an additional indicator, we hence used the ratio of the variance of the controls and the variance of the entire dataset, the fraction of unexplained variance (FUV), which is indicative of the level of noise relative to the biological signal in the data. The FUV overall behaved similarly to the CV and was at minimum at around 20 hours for the uncorrected data. With corrections, this minimal value was largely maintained until the end of the experiment. A lower FUV can be obtained by using maximum slopes rather than individual timepoints.

Although correcting for position and batch effects is essential for high-throughput experiments conducted on agar plates, there is a danger that any normalisation strategy could also create false positives. Specifically, a grid colony positioned next to rapidly growing colony will be smaller (due to nutrient competition) leading to underestimation of the reference surface in that area which will further increase the fitness estimate of neighbouring colonies. This argument applies equally the other way around; cells positioned next to slow growers have better access to nutrients. Indeed, after reference grid normalisation, we often observed a (generally weak but detectable) secondary edge effect for colonies positioned in the next inward row/column (Supplementary Fig. 2D). We found however, that his effect can easily be remedied by an additional row/column median normalisation. To gauge if phenotype exaggeration still presents a significant problem in other parts of the plate, we compared raw and final corrected colony sizes and maximum slopes to the respective sum of all its 8 neighbours. For uncorrected fitness values, there was generally a positive correlation (stronger for colony sizes than for slopes), indicating that regional plate effects dominate over competition between neighboring colonies. This bias was removed after correction. Importantly no negative correlation was observed. We conclude that our grid correction does not lead to any significant effect exaggeration.

### Monitoring cell viability with Phloxine B provides an independent and complementary phenotypic readout to growth assays

The addition of phloxine B to agar medium stains colonies in different shades of red, reflecting the fraction of dead cells, which can provide an additional phenotype readout from the same colony that is used for growth measurements. To investigate how colony size and redness relate, we used the *pyphe* pipeline to characterise 238 *S. pombe* single-gene deletion strains in 70 conditions in biological triplicates (n= 59,350 total colonies profiled including controls but excluding grid colonies, all data in Supplementary Table 1). The two fitness proxies showed little correlation (Pearson r=-0.088) after correction of colony sizes using the grid approach with subsequent row/column normalisation and correction of redness scores by row/column median normalisation only (Fig. 3A). Many mutant-condition pairs showed a strong phenotype in only one of the two read-outs. Noise levels of redness scores were very low (CV=1.04%) and the biological signal strong (FUV=7.83%). We conclude that the phloxine B redness scores provide robust, precise information on mutant fitness, and serves as a largely orthogonal and independent measure compared to the (well correlated) growth rate or colony size measurements.

**Figure 3:**
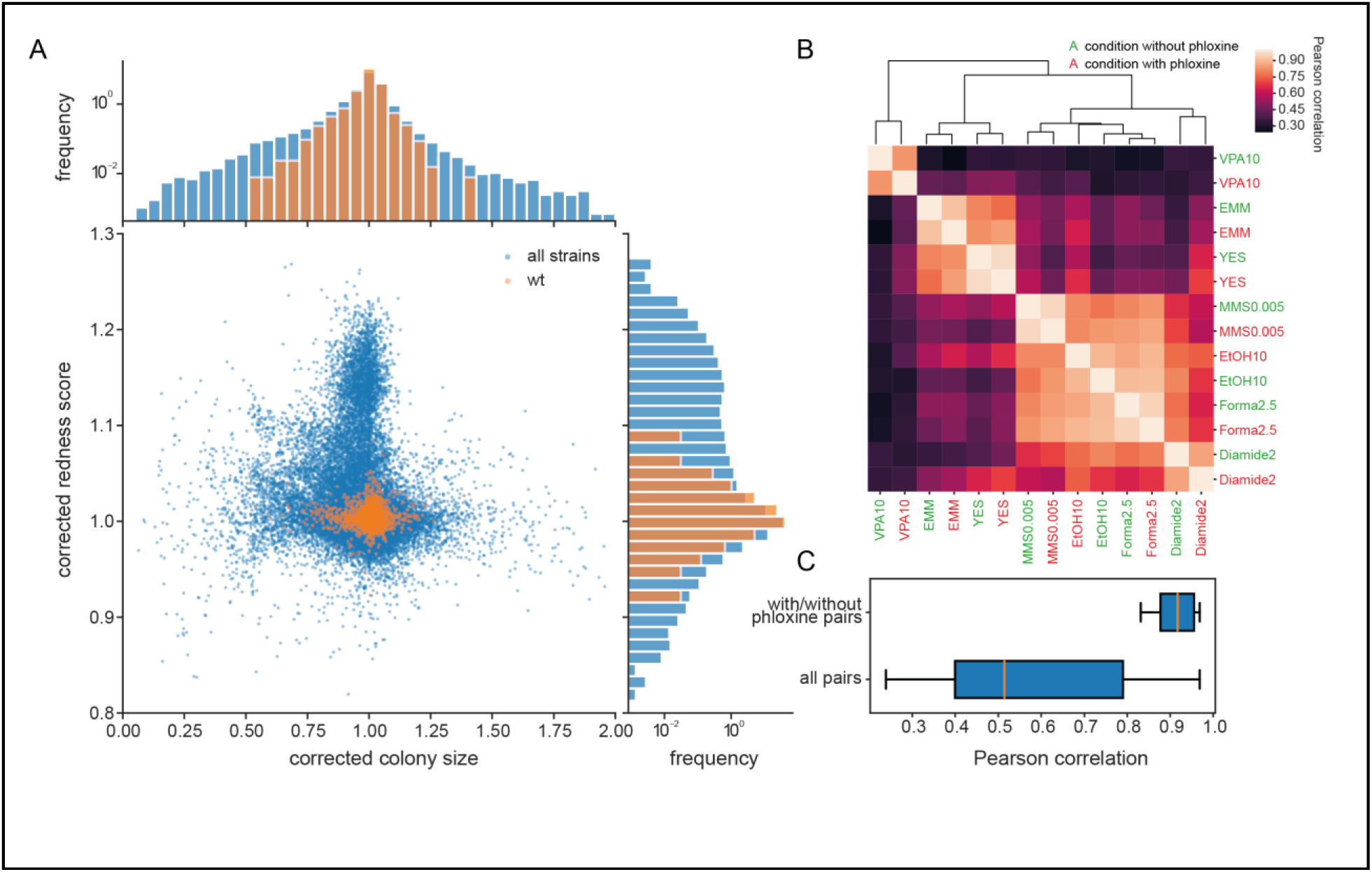
Phloxine B provides an orthogonal, independent fitness proxy. **(A)** Corrected colony size and corrected redness score for 238 single gene knock-outs in 70 conditions (after quality filtering as described in Methods, three biological replicates for each condition-gene pair are shown individually). This plot shows data for individual colonies. The two read-outs are only weakly anti-correlated (r=-0.081) and many mutants-condition pairs show a strong phenotype in only one of the two fitness proxies. Axes were cut to exclude extreme outliers for visualisation purposes. The redness score was robust with a CV of 1.04% and a FUV of 7.83% (histogram on right). For comparison, the CV and FUV of the colony size read-out were 6.1% and 31.5%, respectively (top histogram). **(B)** Clustered pearson correlation matrix of averaged corrected colony sizes (n=3) for 7 conditions with and without phloxine B. Repeats with and without dye consistently cluster together indicating general robustness of our measurements across batches and no substantial mutant-condition-dye interactions. **(C)** Boxplot comparing the pairwise correlation between conditions with and without phloxine B (median=0.92) and all possible pairs from (B) (median=0.51).

Phloxine B can be toxic if exposed to light (Qi et al., 2011), so we tested whether phloxine B changes growth parameters by determining colony sizes for our mutant set in 7 conditions. Measurements with and without phloxine B were performed in different batches and in different weeks to exclude that batch effects drive a closer correlation. Within the 14 phenotype vectors measures in total, identical conditions with and without phloxine clearly and consistently clustered together (Fig. 3B). The median correlation for the 7 condition pairs with and without phloxine was 0.92, which was substantially higher than that of all possible pairs from the 14 phenotypes (Fig, 3C). Overall, this result agrees with a study using rapamycin which has found that phloxine B does not interact with growth in that condition (Lie et al., 2018).

### Phloxine B staining informs about fraction of live cells in colony

Finally, we tested whether and how the colony redness score relates to the viability of cells in the colony. We determined colony composition and viability status at the single cell level using ImageStream flow cytometry. Across 23 samples, obtained from colonies with varying redness scores (Fig. 4A), phloxine B staining classified cells into three populations (Fig. 4B,C): 1) live cells which showed a background level of staining, 2) dead cells which were brightly stained, and 3) lysed or damaged cells which showed no staining. The fraction of live cells (live/dead+lysed) was inversely correlated (Pearson r=-0.88) with colony redness scores obtained with *pyphe-quantify* and row/column median corrected by *pyphe-analyse* (Fig. 4D). This appears paradoxical given that lysed cells are not stained in the ImageStream using a cytometry running buffer, but phloxine B is present in the medium surrounding the colonies where it can enter lysed cells without being washed out. We next asked how well phloxine B staining agrees with a distinct, established dead-cell stain (LIVE/DEAD). In wild-type cells, staining with both dyes agreed closely (accuracy 99.3% using LIVE/DEAD classification as ground truth, Fig. 4E,F). We conclude that phloxine B staining, combined with our imaging and analysis pipeline, provides a sensitive and accurate readout reflecting the proportion of live/dead cells in a colony.

**Figure 4:**
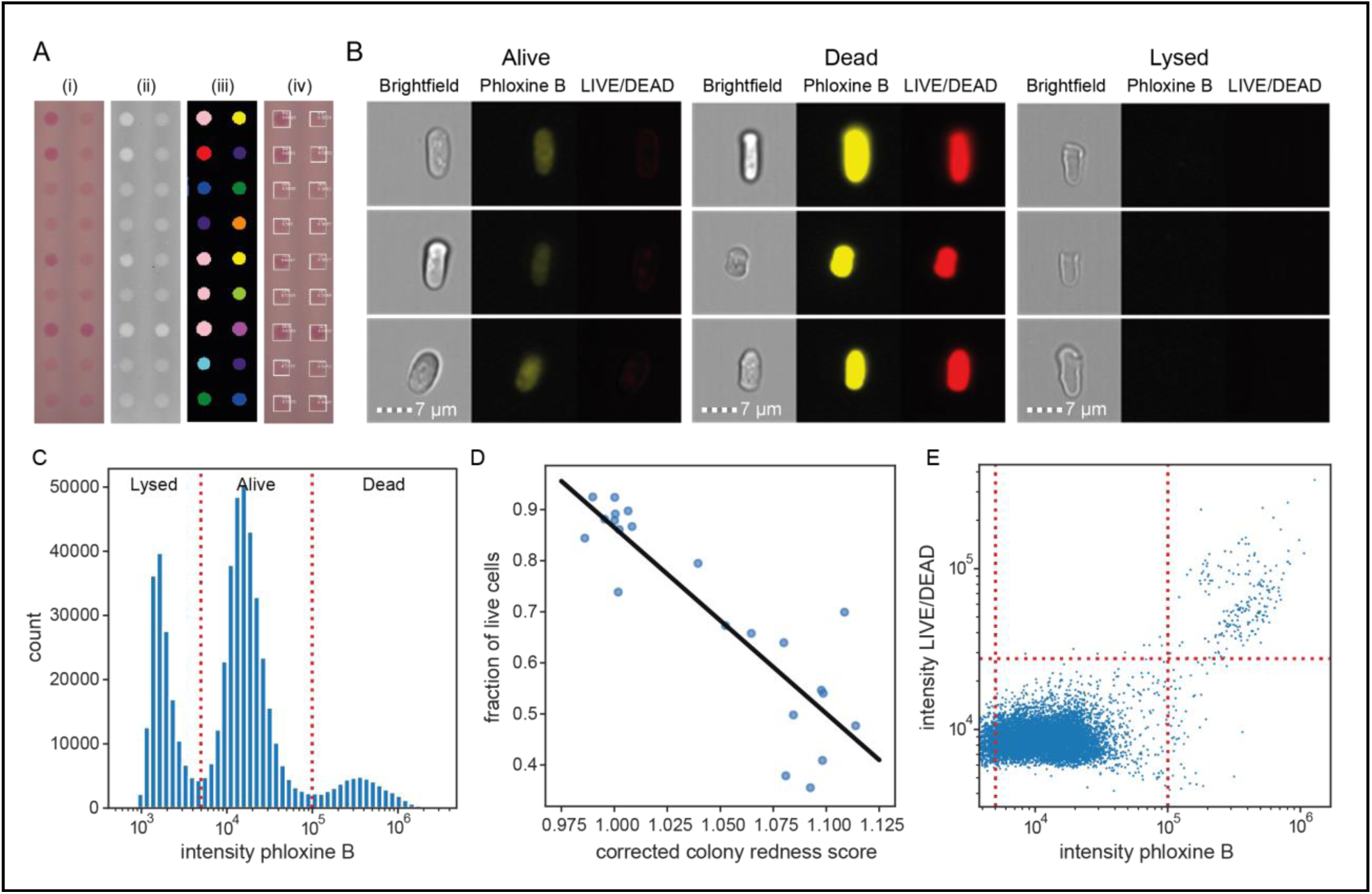
Phloxine B staining reflects percentage of dead cells. **(A)** Example of colony redness score extraction by *pyphe-quantify* in *redness* mode. From the acquired input image (i), colors are enhanced and the background subtracted (ii), colonies are identified by local thresholding (iii), and redness is quantified and annotated in the original image (iv). **(B)** Representative cells for alive, dead and lysed cells using imaging flow cytometry (ImageStream) analysis. Lysed cells show no signal in either the phloxine B or LIVE/DEAD channels. Live cells show an intermediate signal intensity in the phloxine B channel but no LIVE/DEAD signal. Dead cells are brightly stained in both channels. **(C)** Histogram of intensities in phloxine B channel across 23 samples with three populations (lysed, alive and dead) clearly resolved. **(D)** Fraction of live cells (live/(lysed+dead)) by ImageStream correlate with colony redness scores (corrected by row/median column normalisation) obtained with pyphe. **(E)** Co-localisation of phloxine B stain with LIVE/DEAD stain for the standard lab strain *972*. **(F)** Comparison of Phloxine B staining with LIVE/DEAD stain by ImageStream. Both readouts agree with 99.3% accuracy using the illustrated thresholds.

## Conclusion

High-throughput colony-based screening is a powerful tool for microbiological discovery and functional genomics. Using a set of diverse wild yeast strains, we show that the fitness correction approach implemented in *pyphe* effectively reduces noise in the data. Importantly, for endpoint measurements the corrected fitness is independent of the exact time point, as long as a late time point is chosen, and late colony sizes are tightly correlated with maximum slopes of colony areas. This finding has two important implications. First, our results show that growth-rate measurements do not necessarily boost the phenotyping, especially as one can compensate for the reduced precision of end-point measures by measuring more independent replicates. Second, little if anything is gained from precisely pre-defining incubation times of assay plates prior to scanning. Instead, plates can be simply incubated for longer (usually 2-5 days for fission yeast), especially if the assay condition slows down growth.

Furthermore, we show that colony viability measured by phloxine B staining and image quantification by *pyphe* provides a largely orthogonal and independent readout to colony sizes, thus offering an additional trait for mutant profiling. Redness scores obtained with the *pyphe* pipeline closely reflect the number of live cells in the colony. This opens up new avenues of investigations, e.g. for high-throughput chronological lifespan experiments. It will be important to examine the relationship between redness scores and live cells if the proportion of live cells drops to very low levels as the redness signal may saturate. Potentially even more information may be contained in the distribution of dead cells within a colony, which is hard to describe quantitatively and not reported by *pyphe-quantify*.

Here we present the *pyphe* toolbox and underlying python package to assemble a versatile pipeline for analysing fitness-screen data. *Pyphe* is an all-in-one solution enabling image acquisition, quantification, batch and plate bias correction, data reporting and hit calling. *Pyphe* is flexible and accepts growth curves and endpoint measurements as well as colony sizes and staining as input. *Pyphe* functionality is provided in the form of multiple separate, simple and well-documented command line tools operating on human-readable files. *Pyphe* is written for the analysis of extremely large data sets (thousands of plates, millions of colonies), and its modular design allows the easy integration of other, future tools and scripting/automatisation of analysis pipelines which aids reproducibility.

## Software availability statement

*Pyphe* is open software published under a permissive license. We welcome bug reports, feature requests and code contributions through www.github.com/Bahler-Lab/pyphe.

## Materials and Methods

### Wild strain test data set

An overnight liquid culture of strain *972 h-* in YES medium was pinned in 96 (8×12) format on YES agar medium, using a RoToR HDA pinning robot (Singer Instruments) and grown for two days at 32°C. This grid was combined with randomly arranged plates of the 57 wild strains in 1536 (32×48) format and grown for 2 days at 32°C. This plate containing the grid for 972 standard strain lab, every fourth position was copied onto fresh YES agar media, using the 1536 short pinning tool at low pressure. The plate was placed in an EPSON V800 scanner in an incubator at 32°C and images were acquired every 20 minutes for 48h using *pyphe-scan-timecourse*. Growth curves were extracted using *pyphe-quantify* in timecourse mode with the following settings: --s 0.1. Growth curve parameters were extracted with *pyphe-growthcurves*. Individual images were analysed with *gitter* using the following settings: --inverse TRUE -- remove.noise TRUE. Grid correction and subsequent row/column median normalisation of maximum slopes and individual timepoints was performed in *pyphe-analyse*.

### Knock-out test data set

238 mutants, broadly spanning GO Biological Function categories plus several uncharacterised genes, were selected from a prototroph derivative of the Bioneer deletion library (Sideri et al., 2015). Strains were arranged in 384 (16×24) format with a single 96 grid placed in the top left position, so that the grid includes one colony in every four positions within the 384 array. To prepare replicates, this plate was independently pinned 3 times from the cryostock on solid YES media for each batch. From these plates, colonies were then spotted on assay plates containing various toxins, drugs or nutrients. The conditions used in Fig. 3B are: EtOH10 is YES+10% (v/v) ethanol, VPA10 is YES+10mM valproic acid, MMS0.005 is YES+0.005% (v/v) methyl methanesulfonate, Forma2.5 is YES+2.5% (v/v) formamide, Diamide2 is YES+2mM diamide, EMM is standard Edinburgh Minimal Medium, YES is standard Yeast extract with supplements and 3% glucose. Assay plates were usually grown for 2 days at 32°C but this varied according to the strength of the stress slowing the growth of the colonies. After incubation, images were acquired using EPSON V800 scanners and *pyphe-scan* and quantified with *gitter* (see options above) or *pyphe-quantify* in *redness* mode. Grid correction and subsequent row/column median normalisation of maximum slopes and individual timepoints was performed in *pyphe-analyse*. Row/column median normalisation was only applied to redness data plates. For the size data set, 0-sized colonies and colonies with a circularity below 0.85 were set to NA. Plates with a CV >0.2 or FUV >1 were removed as those most likely represent conditions in which the stress was too strong or where technical errors occurred.

### Imaging Flow Cytometry

We picked 23 colonies with varying redness from the collection of 238 *S. pombe* deletion strains grown on solid YES with 5 mg/L phloxine B for 3 days at 32°C and resuspended in 1 mL of water. For analysis of phloxine B staining, 500µL of this cell suspension were centrifuged at 4000g for 2 min, the supernatant was removed and the pellet resuspended in 75µL of PBS. For analysis of phloxine B and LIVE/DEAD co-staining, 500µL of the same suspension were centrifuged at 4000g for 2 min, the supernatant was removed and the pellet resuspended in 300µL of LIVE/DEAD solution (LIVE/DEAD^™^ Fixable Far Red Dead Cell Stain Kit, for 633 or 635 nm excitation, ThermoFisher Scientific, Cat. no. L34974). LIVE/DEAD solution was prepared according to manufacturer instructions (1:1000 dilution in H2O from a stock solution dissolved in 50 uL of DMSO). The pellet was resuspended and incubated for 30 min in the dark. Cells were then spun down and resuspended in 75µL of PBS.

Immediately prior to analysis, samples were sonicated for 20 seconds at 50W (JSP Ultrasonic Cleaner model US21), and transferred to a two-camera ImageStream®X Mk II (ISX MKII) imaging flow cytometer (LUMINEX Corporation, Austin, Texas) for automated sample acquisition and captured using the ISX INSPIRE^™^ data acquisition software. Images of 5000–12,000 single focused cells were acquired at 60x magnification and low flow rates, using the 488 nm excitation laser at 90 mW to capture phloxine B on channel 3; 642 nm excitation laser at 150 mW to capture LIVE/DEAD cells on channel 11; bright field (BF) images were captured on channels 1 and 9, and side scatter (SSC) on channel 6. For co-stained cell analysis, to generate a compensation matrix, cells stained either with phloxine B or with LIVE/DEAD dye individually were captured without brightfield illumination (BF and SSC channels were OFF). The compensation coefficients were calculated automatically using the compensation wizard in the Image Data Exploration and Analysis Software (IDEAS) package (v6.2). Populations of interest (single focused cells) were gated in IDEAS and the features of interest (dye intensities) were then exported for further analysis using Python.

Intensity values were subtracted by their minimum over all samples (which was slightly below zero) and added to 1 prior to log10 transformation. Thresholds for the three populations were set manually based on the intensity histogram across all samples.

## Acknowledgements

Mimoza Hoti helped with the wet lab part of phenotyping the 238 knock-out mutants. This research was funded by Wellcome Trust Senior Investigator Awards to JB [grant number 095598/Z/11/Z] and to MR [grant number 200829/Z/16/Z], as well as a BBSRC Project Grant to J.B. [grant number BB/R009597/1]. This work was supported by the Francis Crick Institute which receives its core funding from Cancer Research UK (FC001134), the UK Medical Research Council (FC001134) and the Wellcome Trust (FC001134).

## Supplementary Text

### Supplementary Text 1 - Image Quantification Algorithms

**Supplementary Figure 1:**
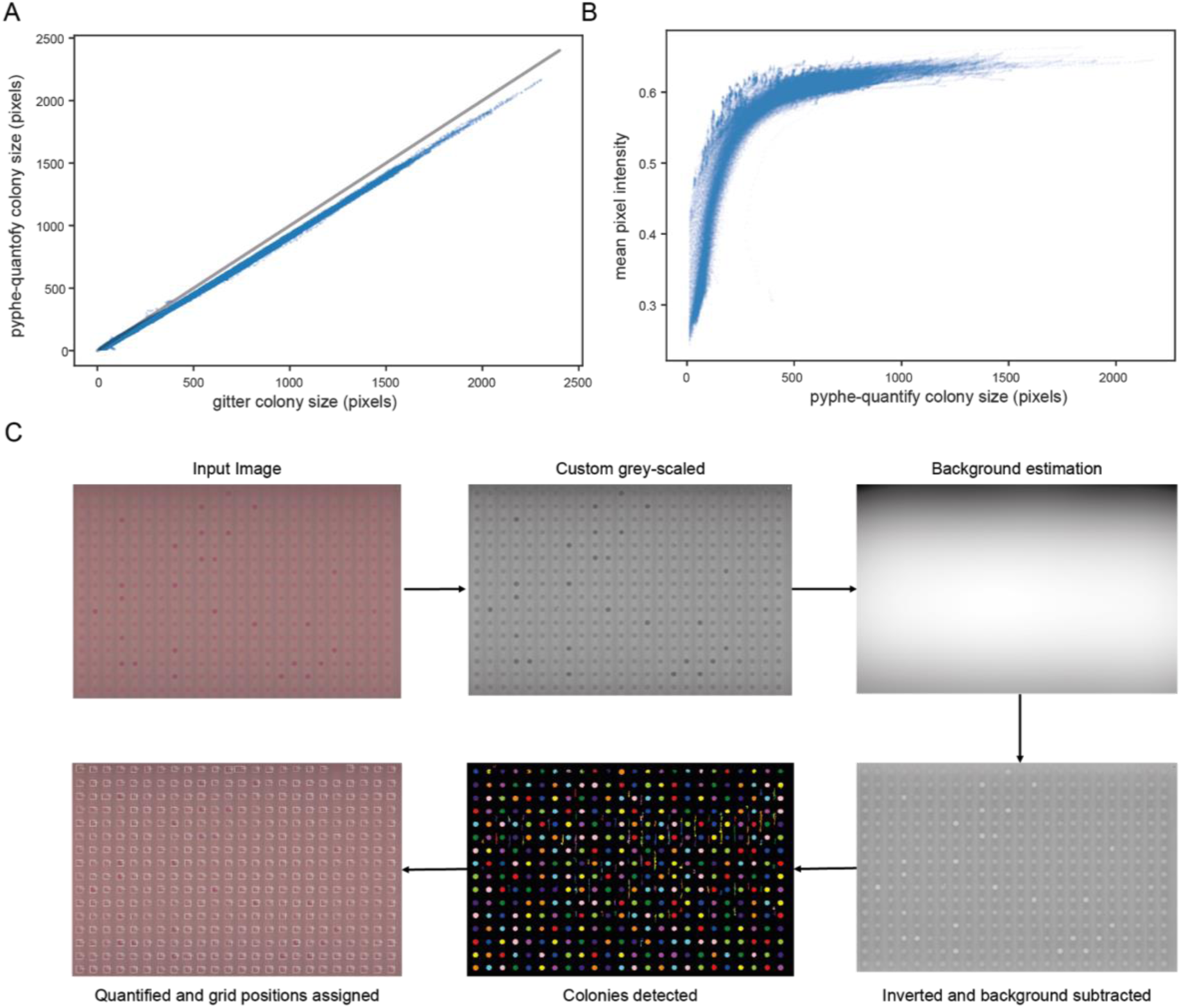
All 144 images from the image time course of 57 wild strains in replicates in 1536 format were analysed *with pyphe-quantify batch* and with *gitter* using remove.noise=TRUE and inverse=TRUE settings. **(A)** Colony area measurements obtained with *pyphe-quantify* are very tightly correlated with those obtained with gitter (r=0.9991). In terms of absolute values, colony size estimates are consistently lower which is due to the default thresholding settings. **(B)** Mean pixel intensities (averaged over all pixels identified as belonging to the same colony) depend strongly on colony size (thickness). This signal quickly saturates as colonies become larger, as previously described by Zackrisson et al. (2016). **(C)** Illustration of image analysis algorithm used by *pyphe-quantify* in redness mode.

*Pyphe-quantify* is a command line tool for the analysis of images containing microbial colonies based on *scikit-image (van der Walt et al., 2014)*. By default, it analyses all .jpg image files in the directory where it is executed, alternatively the user can set a pattern to specify input images. Grid positions are defined by the user in the form of the number of rows, columns and coordinates of the top left and bottom right colonies.

In *batch* mode, *pyphe-quantify* will analyse colony sizes in each image individually. First, morphological components are identified by thresholding the image. By default the Otsu method (Otsu, 1979) is used to find the threshold, but this can be tuned by the user by providing a coefficient to be used with this threshold or an absolute threshold in the config file. Components are then filtered by size to exclude erroneous identification of small particles, such as dust, as colonies. Border components are removed. Components are then matched to grid positions. By default, a component is assigned to a particular position if it is less than a third of the distance between two grid positions away from a position. This threshold can be set by the user. This means that in case of missing colonies, there will be no data for the corresponding grid correction and this position will be missing from the output file (i.e. it will not be 0). Similarly, two components can be assigned to the same grid position in the case of contaminations. This can be disabled by the user to retain only the component nearest to the grid position. When reporting all colonies, *pyphe-quantify* can be used for plates without arrayed colonies. An output table is saved which contains colony area, mean intensity (an estimator that reflects thickness), circularity, perimeter and centroid coordinates. Area measurements are in very close agreement with those obtained with *gitter* (Supplementary Fig. 1A). Mean intensity measurements show no dependence on colony size except for the smallest colonies, indicating that the signal is usually saturated (Supplementary Fig. 1B). *Pyphe-quantify* exports a qc image for every image analysed indicating the identified colonies and their assigned grid positions.

In *timecourse* mode, the final image of the time course is analysed as described above and the obtained mask (indicating the position of each colony) is then applied to all previous images of the time course. The background subtracted mean intensities for all images combined, i.e. the growth curves, is reported in a single file.

Finally, *pyphe-quantify* can analyse colony redness (Supplementary Fig. 1C). Phloxine B stains dead cells within the colonies and these are usually not homogeneously distributed upon close inspection. However, for simplicity, we have developed an image analysis workflow which extracts the mean redness of colonies in high-density arrays, providing a single quantitative readout. We decided to use reflective scanning with our Epson V800 scanners (implemented in *pyphe-scan*). This is fast and produces images with consistent properties, but with the caveat that the focus position is just above the scanner glass and colonies are therefore somewhat out of focus. Additionally, there is a strong, uneven background signal from the media and colour artefacts (appearing as bright stripes between colony columns) which required a different image analysis approach. The images are first adjusted to make colony redness more visible by multiplying the red, green and blue channels by 0, 0.5 and 1, respectively and their sum is taken to produce single-channel/grayscale images. The background value for each pixel is estimated by blurring the image with a Gaussian filter with a standard deviation of the number of pixels in the image divided by 10000. The background is subtracted from the image which is then inverted.

Colonies are then detected by local thresholding and processed further as described for *batch* mode above. The mean intensity for each colony is computed from the processed image and reported in a similar file as described for *batch* mode. The produced QC images allow to verify grid placement and visualise the colour readout on the actual image.

### Supplementary Text 2 - Spatial Correction Algorithms

**Supplementary Figure 2:**
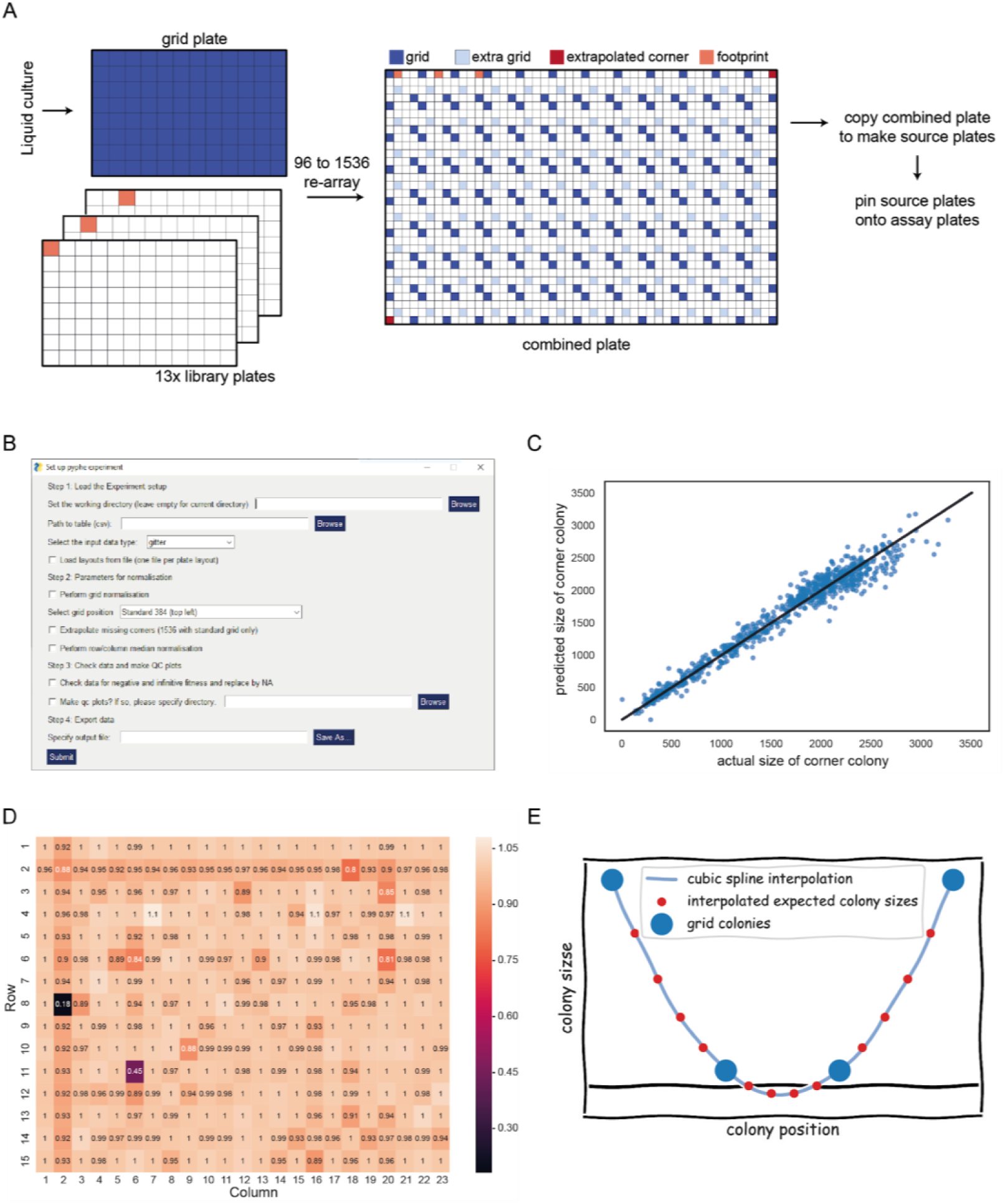
Pyphe normalisation algorithms. **(A)** Placement of two 96 grids in opposite comers of 1536 plate maximises grid coverage and only uses 1 in 8 positions for normalisation purposes. **(B)** GUI for running pyphe. **(C)** Prediction of missing comers based on neighbours. Shown are predicted versus actual colony sizes for top left and bottom right comers for an experiment comprising 375 plates. The linear regression model in this case was y = 1.24 * horizontal neighbour + 0.16 * vertical neighbour - 34. The R^2^ was 0.96. **(D)** Secondary edge effect introduced by grid normalisation. **(E)** Extremely small grid colonies can result in negative values for expected colony sizes.

This text describes the steps performed by *pyphe-analyse* during the analysis of a typical batch of plate images accessed through the GUI (Supplementary Fig. 2B). All functions and objects are also available for use as a python package. *Pyphe* uses the pandas, scipy and numpy packages. At the core of *pyphe* is the Experiment object which is initialised from the Experimental Design Table (EDT). The EDT is first checked for obvious errors, including the uniqueness of plate IDs and paths to data files and if these files exist. Data from the image analysis output files is then loaded using appropriate parsers. Layout files are loaded if set by the user.

Spatial normalisation is then performed if requested by the user. *Pyphe* implements a grid correction procedure similar to that used by (Zackrisson et al., 2016). In that paper, the authors use 1536 format arrays and place a 384 grid in the top left position of the plate. This means one quarter of plate positions are taken up by the grid. It also creates a problem because the right and lower edge of the plate are not covered by the reference grid. We have developed a small improvement of this technique by placing two 96 grids in opposite (top left and bottom right corners, Supplementary Fig. 2A). This leaves only two small corners of the plate (bottom left and top right) not covered by the interpolated grid surface. We solve this by extrapolating the grid by estimating the theoretical colony size of a grid strain in those corners using a linear model and the colony sizes of the two neighbouring grid colonies as input. Model parameters are determined for each experiment based on all plates using regression. We typically achieve accuracies of >90% (Supplementary Fig. 2C). This allows us to use grid correction over the entire plate without loss of data.

We have noticed that doing the grid correction this way slightly over-corrects for the edge effect for strains in the row neighbouring the edge (Supplementary Fig. 2D). This is because the edge effect is usually restricted to the outermost edge only. But the values of the reference grid in the next row/column will be most strongly determined by the grid colony on the edge, leading to an over correction (underestimation of fitness) in these positions. We therefore perform an additional row/column median normalisation after grid correction to remedy this, which is however only possible when the grid is the same as the library background. *Pyphe* offers the option to only perform one or both normalisations.

In some cases, the normalisation procedures can lead to artefacts. Grid normalisation can result in negative corrected fitness values if grid colonies are very small in a region (Supplementary Fig. 2E). Row/column median normalisation can produce infinite values if more than half the colonies in a row/column have size 0. These artefacts are detected by pyphe and set to NA.

*Pyphe* gives the option to produce QC plots for each plate in the experiment in which case a pdf file will be generated containing heat maps for all numerical data associated with that plate. Finally, all data is collated in a single table which contains position information, layout information, all metadata provided by the user in the EDT, raw and corrected fitness values and details about the grid correction.

**Supplementary Figure 3:**
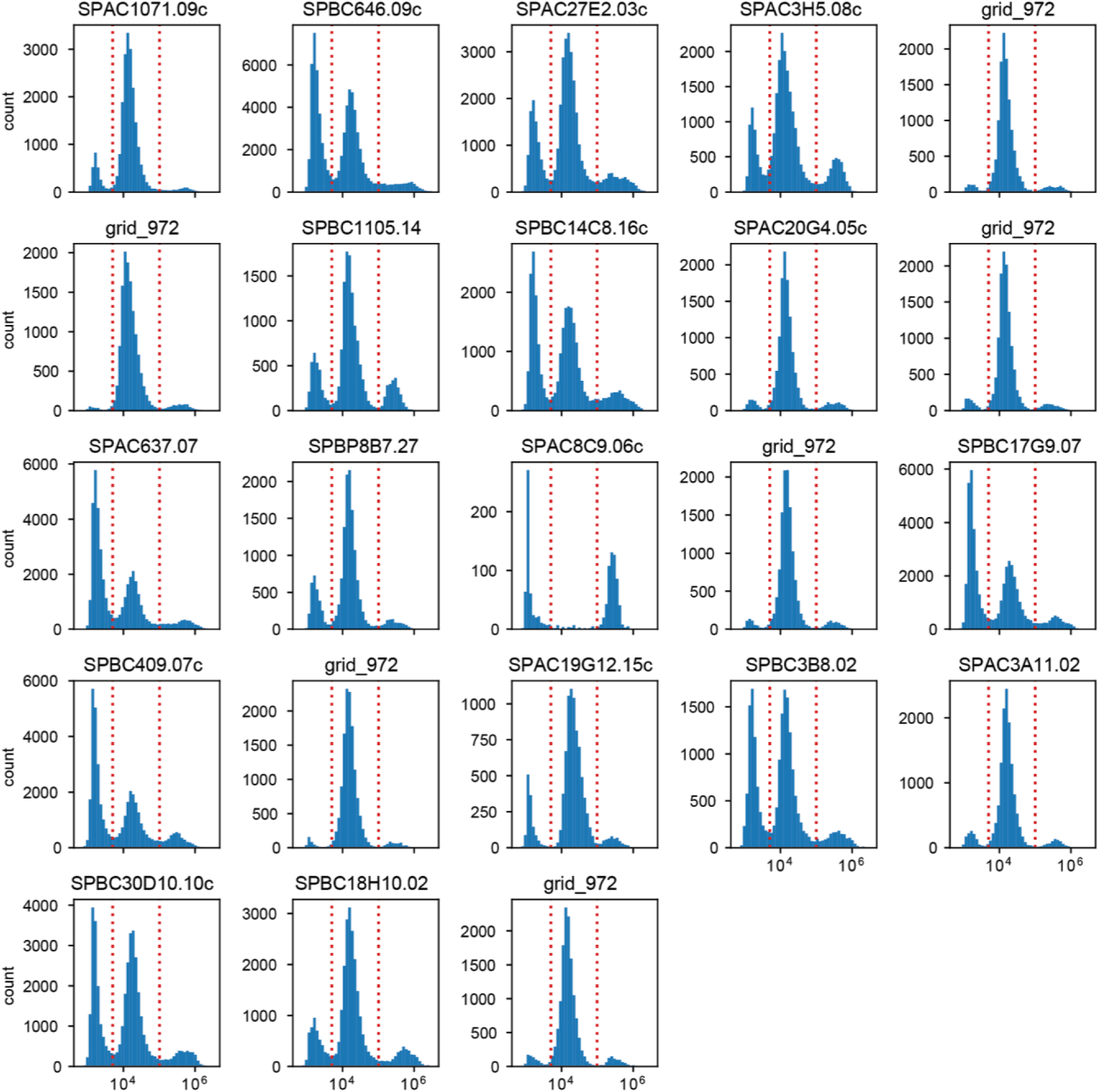
Distribution of phloxine B intensities for different *S. pombe* knock-out mutants obtained by imaging flow cytometry. Plot titles indicate the gene which was knocked out. The standard laboratory strain *972* was measured in 6 biological replicates from samples obtained from different colonies.

### Supplementary Text 3 - Plate Handling Protocol

#### Materials and Reagents

- Sterile yeast medium with and without 2% agar (we preferentially use YES or EMM for *S. pombe*)
- Serological pipette and pipette pump
- Rectangular plates (PlusPlates, Singer Instruments)
- ROTOR pin pads (96 long, 96 short, 384 or 1536 short)
- 96 well sterile plates
- Phloxine B (Merck)

#### Equipment

- Laminar Flow Cabinet
- Microwave oven
- Incubator
- Pinning robot (ROTOR HDA, Singer Instruments or similar)
- Scanner (Epson Perfection V800) connected to Linux computer
- Fixture to hold plates in place on scanner (cutting guide available at www.github.com/Bahler-Lab/pyphe)

#### Procedure

##### Overview

The grid strain is prepared to grow in 96-format plates to make grid plates. Grid plates are combined with library plates to make combined plates. Combined plates are copied onto fresh agar plates to make source plates. Assay plates (containing treatments of interest) are inoculated from source plates. Assay plates are imaged and analysed further.

1. **Plate pouring**
  a. Heat media in the microwave with occasional mixing until completely melted. Let the media cool to approximately 60°C.

> **Warning**: Superheated agar media can pose a serious risk. Proceed carefully, never heat sealed containers and wear appropriate protective equipment.
  b. If drugs are to be added to the media mix them in the media before pouring.
  c. For phloxine B assays, add this reagent at a final concentration of 5mg/L prior to pouring. Note that phloxine B is sensitive to light so it is advisable to store and incubate plates in the dark wherever possible. Phloxine B is also sensitive to oxidising agents and therefore incompatible with such assay conditions.

> **Tip**: A 1000x aqueous stock can be kept in the fridge in the dark for several weeks.
  d. Place plates on a flat surface in the sterile hood and pipette 40 ml of media in each.

> **Tip**: Take up 5ml more than required to avoid bubble formation. If bubbles occur, remove them by sucking them back up into the pipette.
  e. Let plates dry for approximately 30 minutes. Correct dryness is important, if the plates are wet colonies will diffuse into agar.
2. **Plate storage and handling**
  a. Drugless plates made to preserve or wake up collections can be stored in the fridge for a week, but plates should be removed from the fridge and let them acclimate to room temperature prior to any experiment.
  b. Plates containing phloxine B should be stored in the dark, as phloxine B oxidizes in the presence of light.
  c. We recommend preparing assay plates containing drugs on the day of the experiment or the evening prior to the experiment taking place. If this is done, store them appropriately and keep them well wrapped to prevent uneven drying, and upside down to prevent condensation forming on the surface of the media. Let the plates reach room temperature before pinning.
3. **Preparation of source plates: where to locate your grid, controls and your test strains**
  a. **Preparation of the grid plate**
    i. From a cryostock, streak out the strain that will be used as the control ‘grid strain’ on agar media (can be in a conventional Petri dish, add appropriate antibiotics if required). We advise to pick a standard strain which makes the comparative fitness value obtained in the end easily interpretable, e.g. the background strain in case of mutant collections. In general, the grid strain’s fitness should not be extreme (much higher or lower) than the strains to be assayed.
    ii. Grow until colonies suitable for picking have formed (approximately 2 days at 32°C for *S. pombe*).
    iii. Inoculate one colony of the grid strain into 30ml liquid media (e.g. YES in the case of the standard 972 *S. pombe* strain) and grow for ∼24 hours with shaking.
    iv. Pour the grid strain culture into an empty PlusPlate and use the ROTOR to pin onto solid agar media in 96 format using 96 long pin pads.

> **Tip**: Make several copies as needed. You can pin approximately. 10 times from one grid source plate.
    v. Wrap the plates in cling film and place upside-down in incubator to avoid condensation over the colonies. If the plates are not properly wrapped they will dry unevenly on the edges and will not be suitable
    vi. Grow up for approximately 2 days until suitable colonies for pinning have formed.
  b. **Preparation of the library to assay**
    i. **If you are starting from an established library**
      1. If you are using an established yeast library stored in 96- or 384- well format, wake up the library onto the appropriate selective agar media using the ROTOR and let it grow until colonies are visible at the appropriate temperature (32°C for the *S. pombe* Bioneer deletion collection).
      2. Once the colonies are grown refresh the plates onto selective media the same day that you prepare your grid strain plates and let them grow for up to 2 days at the appropriate temperature.
    ii. **If you are arranging your own library**
      1. Prepare fresh colonies of the strains that will be used on solid agar media plates.
      2. Design your library layout. Every plate should contain several wild type controls (at least 10). Plates should contain no or few empty spots, but do include a footprint to mark plate number, orientation and to serve as a negative control. Fill up the rest of the positions with your assay strains and include extra replicates to fill up the plate if required.

> **Tip**: If possible, we recommend to include some positive control strains (that are known to be resistant or sensitive to the stresses to be tested) in the library.
      3. The same day that you will be starting the liquid culture of your grid strain, fill a 96 well plate with the appropriate liquid media. Inoculate each well from a colony, according to your layout.
      4. This 96 well plate can be incubated in a stationary incubator with the lid on for ∼24h at the appropriate temperature.
      5. As with the grid strain plate, use the ROTOR to pin this plate onto solid agar media using 96 long pin pads. Use vigorous mixing (in 3D, 4 cycles) of the source plate.
      6. Wrap the plates in cling film and incubate for 2 days upside down at the appropriate temperature.
  c. **Preparation of the pyphe-ready source plates**
    i. Combine your grid plate and library plates on solid agar medium.
      1. **If your assay plates should be 1536 format**
        a. Please refer to Supplementary Fig. 2A for an illustration of the arrangement process. Using 96 short pin pads and the manual programming mode of the ROTOR, prepare your combined plates by copying the 96 well plate containing the grid strain in the top left and bottom right corner as well as in an additional position in the middle (we normally use the C2 position). Fill the remaining 13 positions with library plates. Record exactly which library plate was used to fill each position and use this information to prepare a layout table of your assay plates.

> **Tip**: This program can be saved and reused.
      2. **If your assay plates should be 384 format**
        a. If you want to work in 384 format, place one grid in the top left corner of each plate. Note that you will lose grid-corrected phenotypes for colonies on the bottom and right edge because these are not covered by the grid. You will also not have a control grid to check the quality of the grid correction. It is usually preferable to use 1536 format with more repeats, even if you have few strains.
    ii. Grow for 1 or 2 days, wrapped in cling film and upside down in an incubator at the appropriate temperature.
    iii. Copy your combined plates onto fresh plates to make your pyphe source plates. This will even out any differences in growth from different inoculum amounts from the previous steps and create a more even spacing of colonies.

> **Tip**: You need to make several copies if you have a large number of assay plates/conditions to be tested. As a rule of thumb, you can use the same plate for pinning ∼6 plates on 1536 format ∼ 8 plates on 384.
    iv. Grow for 1 or 2 days at the appropriate temperature (but keep it consistent). At this stage the plates are ready to be used in the assay.
4. **Phenotyping with pyphe pipeline**
  a. Using the appropriate 384 or 1536 short pin pad, inoculate your source library plates onto your assay plates using the ROTOR. Label your plates clearly with the replicate number, plate layout and condition.

> **Tip**: Use low pressure (around 10% for 384 plates and 4% for 1536 format plates) in order to get a small, consistent inoculum.
>
> **Tip**: Check every time that you did not miss to pin an area of your plate. If this happens, repeat the pinning using a fresh, spare assay plate. If this happens repeatedly, you assay plates were not prepared on a flat, level surface or dried out unevenly.
>
> **Tip**: We recommend using the random offset for picking up the colonies.
  b. Wrap the assay plates on cling film and incubate upside down at the appropriate temperature on an incubator. For 1536 plates and mild stressors around 18 hours incubation is enough time for phenotype observation, 384 format or higher stressors might require further incubation times.
  c. Proceed with image acquisition and data analysis. See manuals and help on github for this. We recommend preparing the Experimental Design Table (which will be later required by *pyphe-analyse*) during scanning, making note of all relevant data and meta-data associated with each plate. The table should contain columns for condition, plate layout, image location, incubation time, batch and scan/pin dates. Save this table in CSV format.

> **Tip**: For large screens containing several batches, consistent naming is essential. We usually define a condition shortcut in a separate table and include the dose without units for brevity, e.g. an entry in the condition column in the EDT may state ‘VPA10’ which is short for YES+10mM valproic acid.
>
> **Tip**: File paths should generally not contain any spaces, non-standard characters or characters forbidden in Unix or Windows file names. Name your condition shortcuts, layouts and replicates accordingly.
>
> **Tip**: Comments or observations which may be important for later analysis (e.g. if there were pinning errors or other issues) should be included in an extra column.
>
> **Tip**: Any additional (meta-)data can and should be included and will be carried through to the data report produced by pyphe.

